# Identification of Influential Variants in Significant Aggregate Rare Variant Tests

**DOI:** 10.1101/2020.10.01.322644

**Authors:** Rachel Z. Blumhagen, David A. Schwartz, Carl D. Langefeld, Tasha E. Fingerlin

## Abstract

**Introduction:** Studies that examine the role of rare variants in both simple and complex disease are increasingly common. Though the usual approach of testing rare variants in aggregate sets is more powerful than testing individual variants, it is of interest to identify the variants that are plausible drivers of the association. We present a novel method for prioritization of rare variants after a significant aggregate test by quantifying the influence of the variant on the aggregate test of association.

**Methods:** In addition to providing a measure used to rank variants, we use outlier detection methods to present the computationally efficient <u>R</u>are Variant Influential <u>F</u>iltering <u>T</u>ool (RIFT) to identify a subset of variants that influence the disease association. We evaluated several outlier detection methods that vary based on the underlying variance measure: interquartile range (Tukey fences), median absolute deviation and standard deviation. We performed 1000 simulations for 50 regions of size 3kb and compared the true and false positive rates. We compared RIFT using the Inner Tukey to two existing methods: adaptive combination of p-values (ADA) and a Bayesian hierarchical model (BeviMed). Finally, we applied this method to data from our targeted resequencing study in idiopathic pulmonary fibrosis (IPF).

**Results:** All outlier detection methods observed higher sensitivity to detect uncommon variants (0.001 < MAF > 0.03) compared to very rare variants (MAF < 0.001). For uncommon variants, RIFT had a lower median false positive rate compared to the ADA. ADA and RIFT had significantly higher true positive rates than that observed for BeviMed. When applied to two regions found previously associated with IPF including 100 rare variants, we identified six polymorphisms with the greatest evidence for influencing the association with IPF.

**Discussion:** In summary, RIFT has a high true positive rate while maintaining a low false positive rate for identifying polymorphisms influencing rare variant association tests. This work provides an approach to obtain greater resolution of the rare variant signals within significant aggregate sets; this information can provide an objective measure to prioritize variants for follow-up experimental studies and insight into the biological pathways involved.

## Introduction

The number of studies that investigate the role of rare genetic variants in complex diseases has been steadily increasing. This is in part due to the hypothesis that rare variant effects might account for the discrepancy between estimated heritability for complex traits and that accounted for by common variant associations [1]. In addition, sequencing technologies have become less expensive such that more studies have the ability to study rare variation in large numbers of individuals or families. A study of circulating adiponectin levels found that among Hispanics, low frequency variants explained more variation in the trait than common variants [2]. Rare variants have been shown to have modest [3] or large effect sizes [1,4]. Rare variants can be associated with increased [5] or decreased risk of disease [6]. Often rare variants influencing a trait are only observed in a few families [7]. Thus, rare variants exhibit a range of association patterns and can exhibit an important contribution to human trait variation.

Studies of rare variation often require either whole genome sequencing or targeted sequencing on large study populations. Even with large sample sizes, there often remains insufficient power for testing rare and uncommon variants individually [8,9]. This is due to the low frequency of observations as well as the higher testing burden, as there are far more rare variants compared to common variants in the genome. To overcome this issue, methods for testing rare/uncommon variants often involve aggregating information across a genomic region [10]. Variants may be grouped into sets based on genomic information (e.g. gene bodies, exons), linkage disequilibrium blocks, association windows, or into overlapping sliding windows of a fixed size in terms of physical distance or number of rare variants included. Methods for testing rare variants in aggregate fall into three main types: burden tests, variance component tests, and combined burden and variance component tests. The underlying assumption of burden tests is that the effects of the rare variants within a given set are in the same direction. Variance component tests ease this constraint and have higher power to detect disease association with a mixture of protective and deleterious rare variants. Given the underlying model by which rare variants are associated with disease in a given region is unknown, tests which combine the burden and variance component approaches weight the given contributions of the burden and variance components to improve power.

Once a set of variants is found to be associated with a phenotype of interest, a logical next step is the identification of the plausible drivers of the association that might be the best statistical candidates for functional studies. Experimental validation of all rare variants in a significant set is generally unreasonable based on time and expense. The ability to narrow down the list of rare variants to those most likely to be contributing to the association signal could help target experimental validation. Additionally, reporting the rare variants most likely to be causal within a significant set of variants focuses functional efforts and can aid in comparison of results across studies.

Several methods have been proposed for statistically identifying the most likely causal variants [11–15]. One method uses a classic backward elimination procedure; a variant is removed from the set if its removal decreases the aggregate test p-value, and this process is repeated until no improvement in the p-value from removing a variant is observed [11]. Related stepwise procedures could be envisioned that use either the p-value or various information criteria (e.g., Bayes information criteria, Akaike information criterion). Another method considers the problem of prioritizing variants in aggregate tests as a variable selection problem using kernel machine methods [16]. Although this method was developed for prioritization of common variants rather than rare variants, conceptually it is reasonable for rare variants. There are also previously published methods which test each variant individually, such as using a Fisher’s Exact Test and applying the adaptive combination of p-values procedure (referred to as ADA) to the resulting p-values [12]. This method has been shown to be more powerful than the backward elimination method proposed by Ionita-Laza, et. al., 2014. Most recently, the BeviMed method applies a Bayesian hierarchical model and makes inference on whether a rare variant is causal based on power posteriors [15]. Methods like the backward elimination method and adaptive combination of p-values are iterative in nature and therefore computationally intensive. In addition, the Bayesian approach is developed for genome-wide testing of rare variant associations in rare Mendelian disorders under specific (unknown) modes of inheritance rather than being more broadly applicable.

We present a novel method for prioritization of rare variants within a given set of variants after the set of variants is found to be significant using aggregate testing methods. Building on the rich outlier-detection statistical literature, we present a computationally efficient approach to be applied following identification of a set of variants that is agnostic to putative function. Our approach, which we refer to as RIFT for <u>R</u>are Variant Influential <u>F</u>iltering <u>T</u>ool, leverages the influence of the variant on the aggregate test of association by quantifying the change in the aggregate test when that variant is removed. It is particularly well suited for rare and uncommon variants, the most common applications of aggregate tests, but is applicable to aggregate testing of variants of all frequencies. When applied to a significant set of rare/uncommon variants, RIFT provides a scheme for quantifying the contribution of an individual variant to the overall association signal, while adjusting for covariates. This method also provides a quantitative measure by which to rank variants for further investigation and several visualizations to aid in evaluation of a region of interest.

## Methods

### Overview

We present the <u>R</u>are Variant Influential <u>F</u>iltering <u>T</u>ool (RIFT) to quantify the effect of each variant on an aggregate test of association (Figure 1). In our simulations, we consider variants with a minor allele frequency (MAF) of <3% (i.e., rare and uncommon), but note that the method is directly applicable to any set of variants, including common variants or mixtures of rare and common variants. First, we describe our jackknife (leave-one-out) approach to obtain a score for each variant when applied to a set of rare variants from an aggregate test. We then describe several outlier detection methods that aim to identify influential variants (IVs) when applied to the variant scores within a set. We perform simulations to evaluate the jackknife scores and the ability of the outlier detection methods to identify variants simulated to be associated with the outcome. Finally, we apply RIFT to recently published rare variant regions found to be significantly associated with idiopathic pulmonary fibrosis [17].

**Figure 1.**
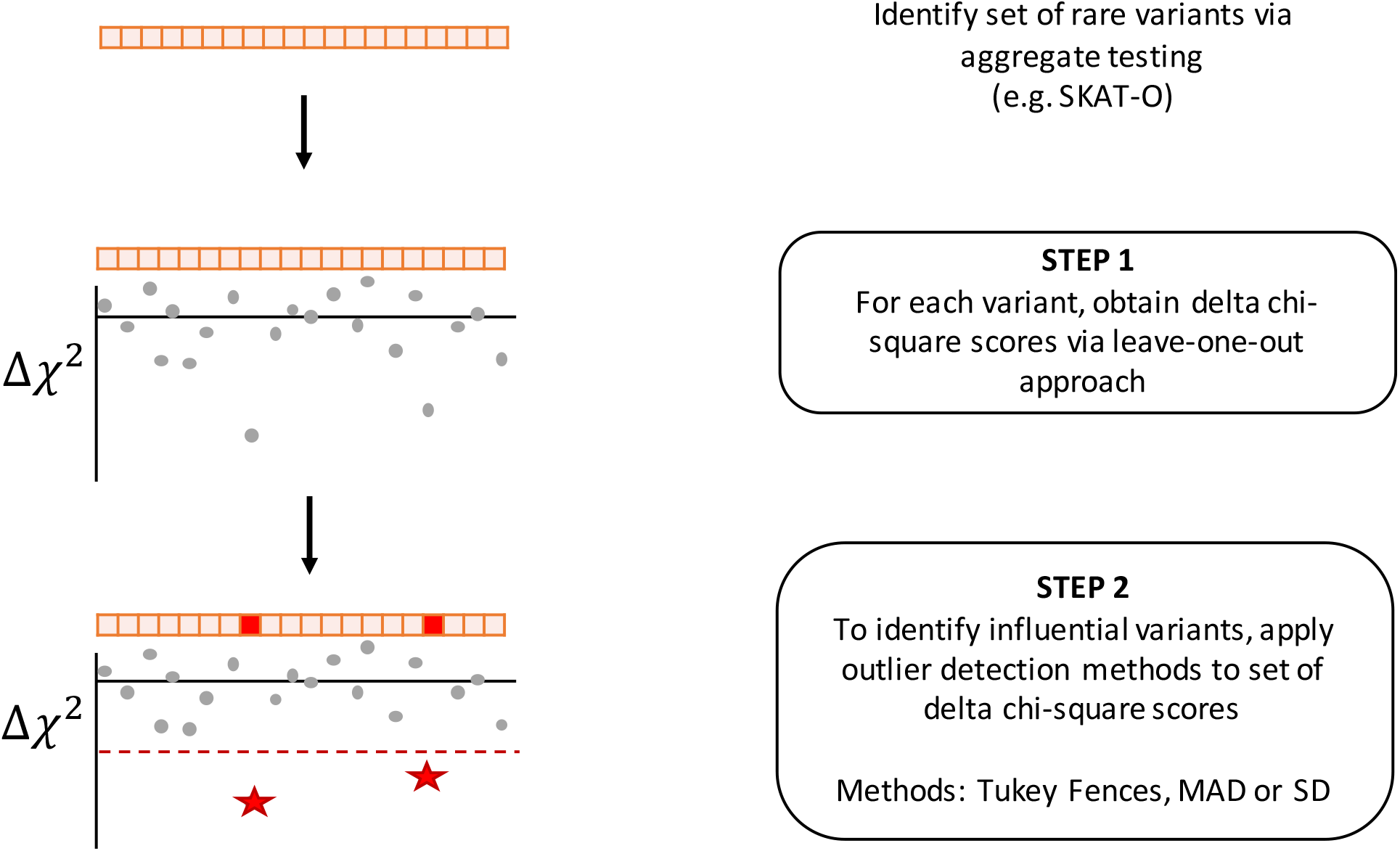
Flow chart of Rift.

### Localization Approach

For a given set of rare variants, we define the p-value resulting from the aggregate test as *p*_(0)_. For rare variant *j*, we calculate the p-value from the aggregate test of the variants within the set excluding variant *j* and refer to this p-value as *p*_(*-j*)_. P-values are transformed into chi-square statistics (df = 1) using the inverse cumulative distribution function (CDF) of the chi-square distribution (Equation 1), where 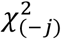 corresponds to *p*_(*-j*)_. and 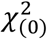. corresponds to to *p*_(*o*)_. For each rare variant *j*, we calculate a delta chi-square score, denoted 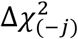, which represents the change of the chi-square statistic when the rare variant *j* is removed (Equation 2). For variant *j*, 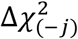 provides a relative measure of how the results of the aggregate test compare with and without variant *j*. Larger negative values indicate larger contributions to the overall test statistic.

Equation 1: Inverse CDF function of the chi-square distribution (df = 1) can be written in terms of the inverse CDF of the normal distribution, *N*(0,1) denoted as *Φ*^-1^.

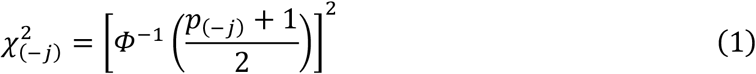

Equation 2: Delta chi-square score

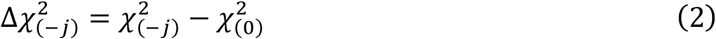

### Outlier Detection for Identifying IVs

The delta chi-square score provides a quantitative measure to rank the rare variants within a significant set according to the impact of that variant on the aggregate test of association. In addition to ranking, it is also desirable to identify the subset of polymorphisms influencing the set’s statistical association. We considered this as an outlier detection objective, whereby unusually large delta chi-square scores across a significant set of variants correspond to the set of association influencing polymorphisms. Sample values more extreme than a pre-specified cutoff are determined to be outliers. Outlier detection methods are rooted in using an estimate of the sample variance that is robust to outliers, and robust measures of spread are often applied for that purpose. We considered two non-parametric approaches to identifying outliers and compared these to a parametric approach.

### Non-parametric variance estimation approaches

In 1977, Tukey defined the commonly-known descriptive univariate boxplot that displays the interquartile range (IQR) as a measure of spread (IQR = distance between the first [Q1] and third [Q3] quartiles; [18]. Boundaries based on the IQR are referred to as “fences”, and observations lying outside the fences are considered outliers. The inner fence is defined by Q1 - 1.5*IQR and Q3 + 1.5*IQR; the outer fence is defined by Q1 - 3*IQR and Q3 + 3*IQR. Note that the outer fence boundary is further away from the median than that of the inner fence boundary, and is therefore more conservative in classification of outliers. The IQR is robust to extreme values in the data and therefore identification of outliers using the IQR are superior to methods that rely on parametric variance estimators in the presence of outliers [19]. Additionally, the IQR and corresponding Tukey fences do not make any distributional assumptions and have been shown to be effective as long as the data are not highly skewed [19].

Another measure of spread that does not assume a parametric distribution for the data is the median absolute deviation (MAD). Identification of outliers based on the MAD is provided in Equation 3 (further details can be found in Leys, Ley, Klein, Bernard, & Licata, 2013). Similar to the IQR, the MAD as a measure of spread is robust to extreme values compared to the standard deviation, which is greatly influenced by extreme values. The MAD requires specification of a constant and a cutoff; as others have used for criteria in outlier detection, we set the constant corresponding to the normal approximation (b = 1.4826) and a cutoff value of three; observations more than three MAD away from the median are considered outliers [21,22].

Equation 3:

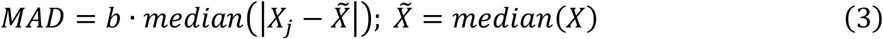

For completeness, we compare the cutoffs based on both the Inner and Outer Tukey fences and MAD to that of the standard deviation (SD), in which observations more extreme than three standard deviations from the mean are considered outliers. This method is often referred to as the three sigma rule [23]. Given that we expect causal variants to reduce the chi-square statistic when removed, we further limited classification of IVs to variants having a negative delta chi-square score.

### Simulated Data

We follow a previously developed simulation approach to generate rare variant data with a binary outcome [24,25]. Specifically, to generate case-control rare variant data under both null and alternative hypotheses, we simulated 10,000 haplotypes for a 1MB genomic region under a coalescent model with parameters consistent with a European population using the software package COSI [26]. We considered 50 different genomic regions of size 3kb and included only rare variants with MAF < 0.03. To generate samples of cases and controls from the population, we repeatedly selected two haplotypes at random and converted these haplotype data to genotypes at each variant location. We determined the probability of subject *k* being a case, defined as *p*_*k*_, based on a logistic regression equation where *β*_*k*_ represents the coefficient for variant *k* (Equation 4). *G*_*jk*_ corresponds to number of minor alleles carried by subject *k* for variant *j* and *β*_*0*_ the disease prevalence. For all simulations, the disease prevalence was fixed at 0.05. Case status, *Y*_*k*_, is specified using the Bernoulli (*p*_*k*_) distribution.

Equation 4:

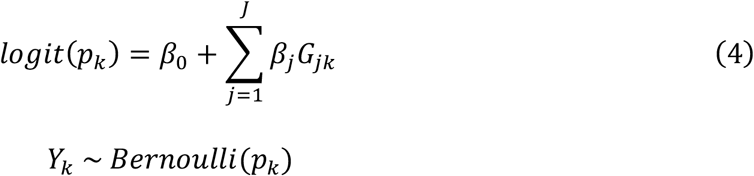

Consistent with previously published simulation of rare variant data, we defined the coefficient for variants under the alternative (Equation 5) to be a function of the population minor allele frequency (*MAF*_*j*_) for variant *j* [24,25]. This relationship between the coefficient and the population MAF results in larger odds ratios for more rare variants. The constant, *c*, defines the strength of association between the causal variants and the outcome.

Equation 5:

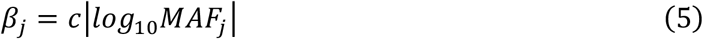

With *c* set to 0.4, this corresponds to an odds ratio of 3.32 for a variant with a MAF of 0.001 and 7.39 for a variant with a MAF of 0.00001. After determining case status for a large sample of individuals, we sub-sampled to obtain a specified number of cases and controls. As expected, the sub-sampling of the population results in some variants (both under the null and alternative) to be no longer observed in a given sample.

Note that our approach can be applied to any such aggregate test (for a review of methods for rare variant aggregate tests, please see Seunggeung Lee, Abecasis, Boehnke, & Lin, 2014). For exposition of the method, we use a combined burden and variance component test, the Sequence Kernel Association Optimal Unified Test (SKAT-O), due to its popularity in rare variant analyses [25]. We used the recommended settings – including a linear weighted kernel, estimation of p-values using the “davies” method and the variant weights as a function of the MAF using the Beta(1,25) distribution. If the SKAT-O p-value met the significance level (here, alpha level of 0.05), we then applied our localization method to obtain a chi-square statistic and delta chi-square score for each variant. We considered the proportion of the total 1000 samples with a SKAT-O p-value that met the significance as an estimate of the power of SKAT-O for that region. SKAT-O performs a grid search to determine the optimal value of the parameter *ρ* that weights the relative influence of the burden and variance statistic. The parameter *ρ* determined to be optimal for the full data was then fixed when calculating the leave-one-out p-values. This insured that within a significant set, the jackknife p-values and corresponding chi-square statistics are comparable. To identify IVs, we applied the previously described outlier detection methods to the resulting delta chi-square scores. For every variant, we calculated the proportion of times it was labelled as an IV across the samples it was observed; for variants under the alternative, this corresponds to a true positive rate and for variants under the null, this corresponds to a false positive rate.

### Comparison with existing methods

We compared the performance of RIFT to that of ADA and BeviMed by applying both methods to simulated datasets. Due to the superior performance and computational efficiency of ADA compared to the backward selection procedure, we chose to limit our comparisons to these two methods [12]. We followed our simulation approach above, where we simulated 50 genomic regions of size 3kb with 10% of rare variants under the alternative and effect size parameter of 0.4. Given RIFT is developed for regions which are previously found to be associated via an aggregate test of association, we restricted our comparisons of methods to regions where the SKAT-O was significant at 0.05. We performed ADA using the default parameters: 1) a MAF threshold of 0.05, 2) calculation of p-values using Fisher’s exact test, 3) assuming an additive model and 4) 1000 permutations for calculation of the final region ADA test p-value. For each replicate, the final set of variants (per-site p-value smaller than the optimal value) were classified as IVs. We applied BeviMed using the default parameters: 1) for each variant, a prior probability of association of 0.01 and b) prior probability of a dominant model of inheritance of 0.5. BeviMed returns the posterior probability of association for both the dominant and recessive models of inheritance. For each of these models, we classified any variant with a posterior probability greater than 0.9 as a IV. We compared the above methods to our localization method (RIFT) using both the Inner Tukey and MAD criteria applied to the delta chi-square scores (referred to as RIFT:Inner Tukey and RIFT:MAD). To compare the performance of the above methods, we calculated the true positive rate and false positive rate across samples in which a variant was observed.

### Impact of Varying Sample Characteristics on RIFT Performance

To determine the robustness of RIFT performance to varying characteristics of a given sample, we performed additional simulations that varied the following parameters: # haplotypes simulated (1,000 and 100,000), sample size (5,000 cases and 5,000 controls), region size (0.75kb), disease prevalence (10%), coefficient of disease association (*c* = 0.8) and proportion of alternative variants (20%). For each, we simulated 500 samples for each of 10 regions while fixing the other simulation parameters to those used in the comparisons with the ADA and BeviMed methods. We summarized these different conditions by the structure of the resulting genotype data (number of variants, etc.), number of samples meeting SKAT-O significance level (power of SKAT-O) and finally in the performance of RIFT such as the relationship between delta chi-square score, MAF and true positive rate (Supplemental Table 1).

**Table 1.**
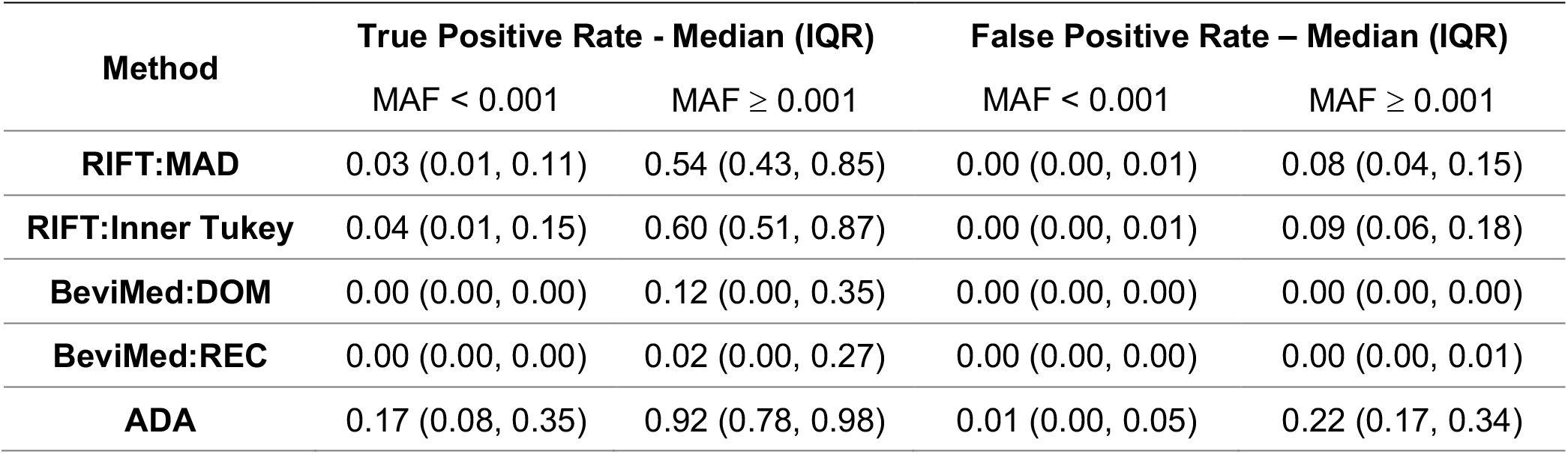
Summary of true positive and false positive rates, stratified by MAF for each localization method. The outlier detection methods for RIFT (MAD and Inner Tukey) were applied to the delta chi-square scores. Simulation included 1000 samples per region, across 50 regions. Data were simulated to have 10% variants under the alternative with effect size parameter *c* = 0.4 (see Equation 5).

### Application to Rare Variant Regions Associated with IPF

We applied our leave-one-out localization method to data from a recently published targeted resequencing study in idiopathic pulmonary fibrosis (IPF) [27] as part of the Global IPF Collaborative Network (http://www.ucdenver.edu/academics/colleges/medicalschool/departments/medicine/GlobalIPF/Pages/GlobalIPF.aspx). We applied RIFT with the goal of identifying rare variants within regions associated with IPF to focus follow-up experimental validation studies. We report results for two regions, each containing 50 rare variants. To provide insight into putative function, we include functional annotation information obtained from SNPDOC (https://wakegen.phs.wakehealth.edu/public/snpdoc3/index.cfm) and HaploReg v4.1 [28].

## Results

### Characteristics of simulated regions

Among the 50 simulated 3kb regions, each region included between 43 and 73 rare variants (MAF < 3%) with a median of 58.5 variants. Though we explored a range of the proportion of variants assumed to be under the alternative, we report the results for simulations where 10% of the variants in a given region were simulated to be under the alternative. Results are qualitatively very similar for higher proportions of variants under the alternative (Supplemental Table 2). After drawing random samples and selecting 1000 cases and 1000 controls to reflect the sampling process, the average proportion of variants under the alternative across the 50 regions ranged from 9.7% to 15.2% with a median of 12.3%. Samples for a given region often contained higher than 10% alternative variants due to the over-sampling of cases to obtain an equal number of cases and controls (Supplemental Table 2). We applied our localization method to samples that had a SKAT-O p-value less than 0.05. With the low proportion of variants (10%) simulated under the alternative and effect size parameter, *c*, of 0.4, the observed median power of SKAT-O across the 1000 simulated samples was 22.0%. As expected, increasing the effect size parameter to 0.8 for the same 50 regions dramatically increased the median power of SKAT-O to 99.1% (Supplemental Table 2).

**Table 2.** Influential Variants identified in two rare variant loci previously found associated with IPF

| Chr:Pos<sup>a</sup> | MA<sup>b</sup> | MAF<sup>c</sup> | Delta Chi-Square | rs# | SNP-DOC annotation | Nearest Gene | Publications/Clinical Notes |
| --- | --- | --- | --- | --- | --- | --- | --- |
| chr5:1294704 | A | 0.00056 | -3.65 |  | coding-synon,ncRNA | TERT |  |
| chr5:1295161 | G | 0.00353 | -17.50 | rs878855297 | ncRNA,untranslated-5 | TERT | - Familial and sporadic melanoma (Horn et al., 2013)<br>- High penetrance, early onset melanoma (Harland et al., 2016)<br>- Cancer cell lines, found to abrogate telomerase silencing and promote tumorigenesis (Chiba et al., 2015) |
| chr5:1295228 | A | 0.00705 | -28.68 |  | unknown | TERT | - Familial and sporadic melanoma (Horn et al., 2013; Huang et al., 2013)<br>- Cancer cell lines, found to abrogate telomerase silencing and promote tumorigenesis (Chiba et al., 2015)<br>- Thyroid cancer (Liu and Xing, 2016) |
| chr20:62324391 | A | 0.05008 | -6.77 | rs41308092<sup>d</sup> | intron | RTEL |  |
| chr20:62324564 | T | 0.00197 | -3.25 | rs398123017 | ncRNA,nonsense | RTEL | - Dyskeratosis congenital (Ballew et al., 2013; Walne et al., 2013)<br>- Hoyeraal-Hreidarsson syndrome (Deng et al., 2013)<br>- Familial interstitial pneumonia (Cogan et al., 2015)<br>- Idiopathic pulmonary fibrosis (Todd et al., 2017) |
| chr20:62324600 | T | 0.00098 | -2.60 | rs373740199 | coding-synon,ncRNA,nonsense | RTEL | - Dyskeratosis congenital (Ballew et al., 2013)<br>- Hoyeraal-Hreidarsson syndrome (Moriya et al., 2016)<br>- Idiopathic pulmonary fibrosis (Todd et al., 2017)<br>- Early-onset Inflammatory Bowel Disease (Petersen et al., 2017) |

<sup>a</sup>NCBI Build 37 coordinates
<sup>b</sup>Minor allele observed in our data
<sup>c</sup>Minor allele frequency observed in our data
<sup>d</sup>HaploReg results: promoter histone marks found in skin, GI and enhancer histone marks found in 8 tissues including lung and a lung carcinoma cell line

### Performance of Our Localization Methods

Across all regions, we found an interesting relationship between the delta chi-square score and the MAF (Figure 2). Delta chi-square scores were more extreme (and more variable) for variants with smaller odds ratios and correspondingly higher MAFs. The delta chi-square score is most sensitive to uncommon variants (0.03 > MAF ≥ 0.001) with modest effect sizes and least sensitive to rare variants (MAF < 0.001) with stronger effect sizes. Positive delta chi-square scores were observed for some variants under the alternative; however, the average delta chi-square score was negative for all but three variants out of a total of 288 variants simulated under the alternative across the 50 regions. The general directionality of the delta chi-square score is consistent with what we expect for causal variants, where the removal of a causal variant results in a larger p-value, smaller chi-square statistic and therefore a negative delta chi-square score. The relationship between the delta chi-square score and MAF was consistent when varying the number of haplotypes simulated; however, we observed an even stronger relationship when increasing the sample size from 1,000 cases and controls to 5,000 cases and controls (Supplemental Figure 2).

**Figure 2.**
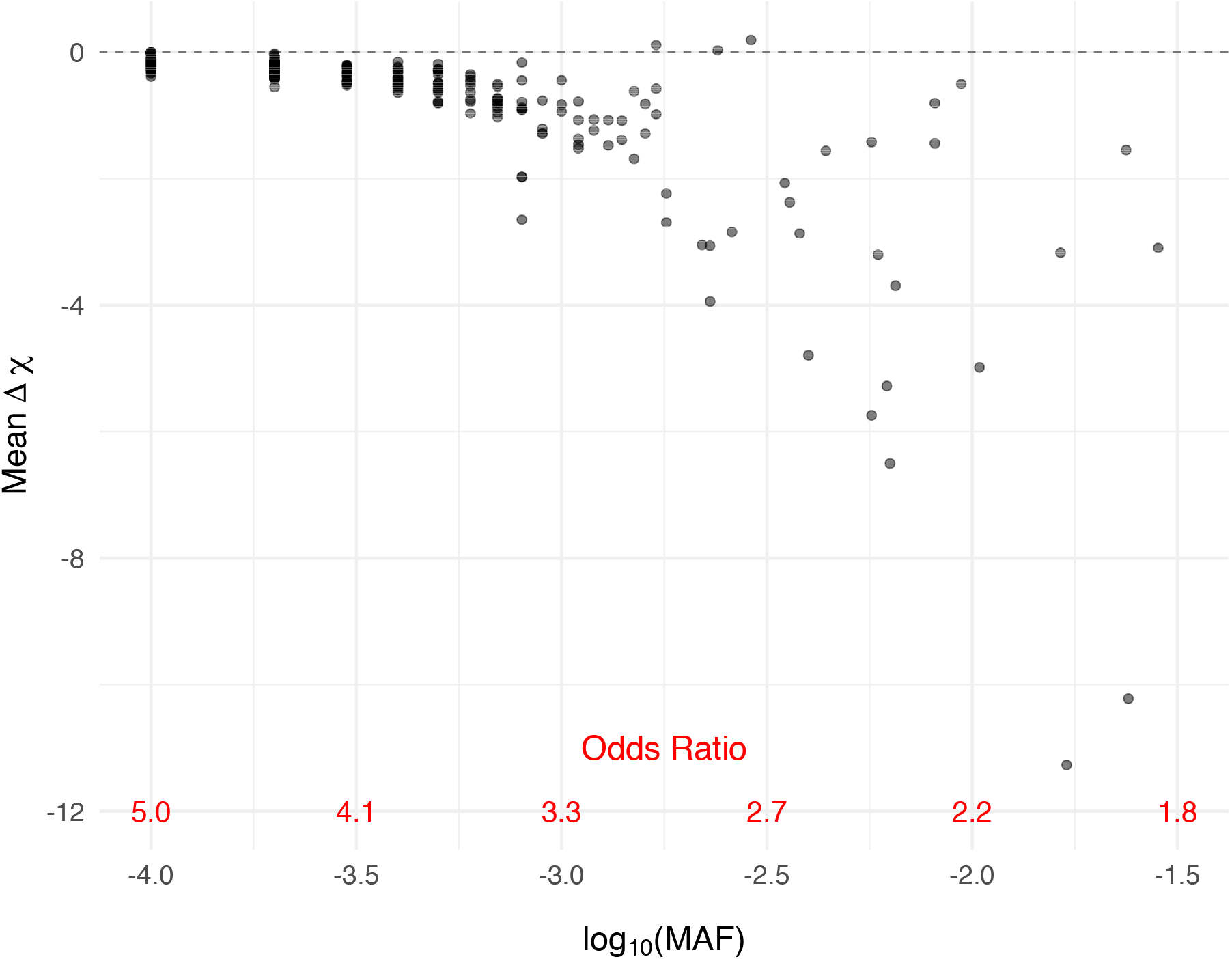
Delta chi-square score more sensitive to the more frequently observed rare variants with correspondingly smaller effect sizes. Mean delta chi-square score plotted by odds ratio for variants simulated under the alternative across all 50 regions. Data were simulated to have 10% variants under the alternative with effect size parameter *c* = 0.4 (see Equation 5).

Consistent with what was observed for the delta chi-square score, all outlier detection methods observed higher sensitivity to detect uncommon variants (0.03 > MAF ≥ 0.001) compared to very rare variants (MAF < 0.001). The Inner Tukey fence had the highest true positive rate (correctly labeling a variant under the alternative as an IV) compared to other methods (Figure 3). For variants with a MAF ≥ 0.001, corresponding to an odds ratio less than 3.32, the Inner Tukey observed a median true positive rate of 0.60 (IQR: 0.51, 0.87) compared to the SD method having a median of 0.30 (IQR: 0.10, 0.59; Supplemental Table 4). Among the 288 variants under the alternative, the Inner Tukey fence obtained the highest true positive rate 94.1% of the time and obtained a higher true positive rate than the other three methods 74.7% of the time. The false positive rate (inaccurately labeling a null variant as an IV) was remarkably low across all four methods, with the SD method having the lowest rate. For variants with MAF ≥ 0.001, the SD method had a median false positive rate of 0 (IQR: 0, 0) and the Inner Tukey a median of 0.09 (IQR: 0.06, 0.18; Supplemental Figure 1, Supplemental Table 4). Taken together, the Inner Tukey has the best characteristics for correctly labeling a variant as an IV, as evidenced by the high true positive rate and low false positive rate. The SD method was substantially more conservative in labeling variants as IVs under the alternative, especially for those variants having a MAF ≥ 0.0001.

**Figure 3.**
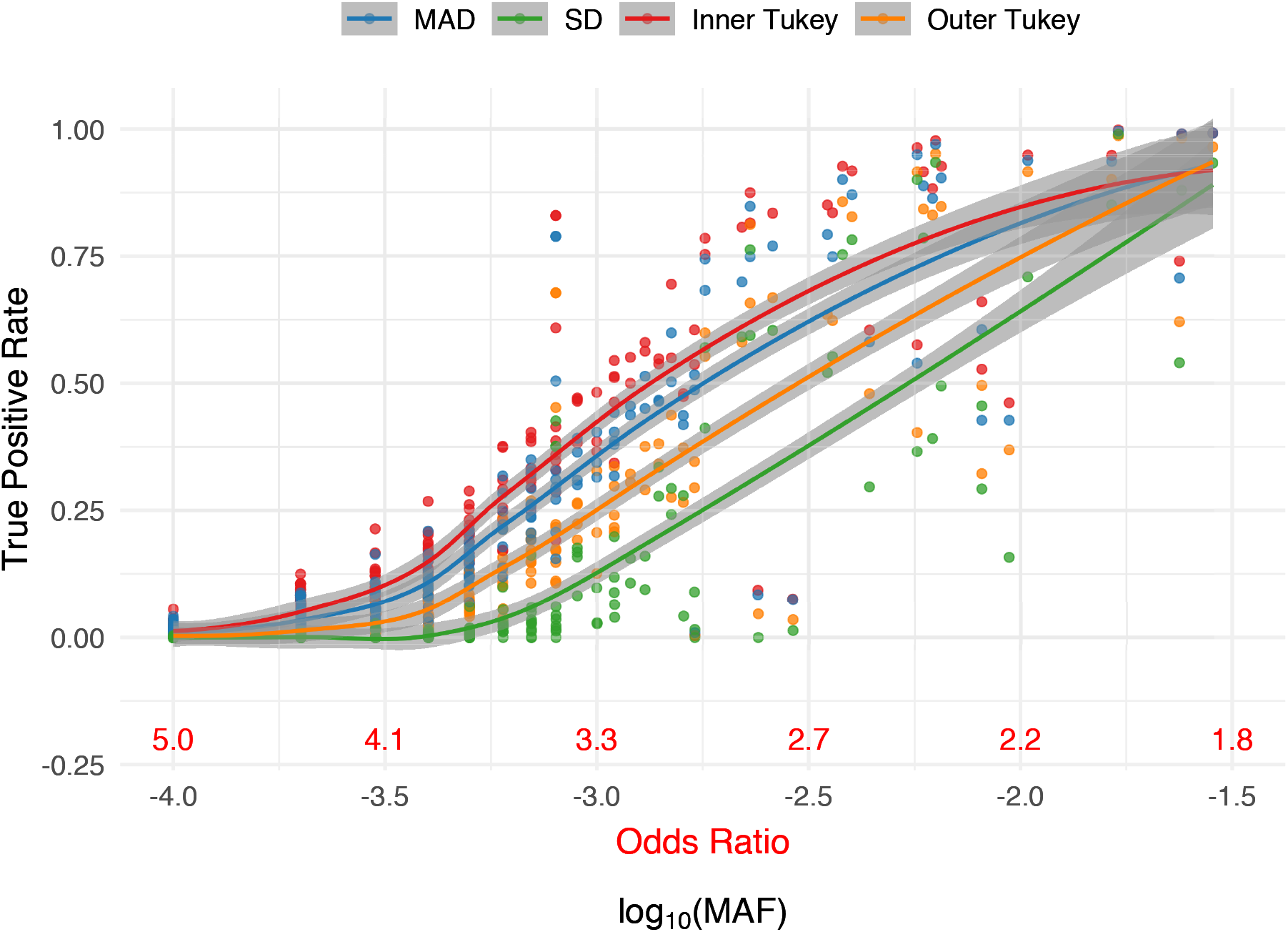
True positive rate (proportion correctly labeled IV) for variants under the alternative for each outlier detection method of RIFT as a function of MAF. Corresponding odds ratio is also provided for reference. Smoothed line and confidence band provided by the loess method. Data were simulated to have 10% variants under the alternative with effect size parameter *c* = 0.4 (see Equation 5).

### Comparison with existing methods

Similar to RIFT, we observed a trend among existing localization methods in terms of having increased ability to identify IVs at uncommon allele frequencies compared to rare. (Figure 4, Table 1). While comparing the ADA and the BeviMed to the Inner Tukey and MAD, the ADA had the highest true positive rate across the entire MAF spectrum we considered, whereas BeviMed (dominant and recessive) observed the lowest true positive rate. For uncommon variants (MAF ≥ 0.001), the ADA had a median true positive rate of 0.92 (IQR: 0.78, 0.98) compared to BeviMed under the dominant model having a median of 0.12 (IQR: 0.00, 0.35). The higher sensitivity of the ADA to correctly label IVs for rare MAFs is at the cost of having a high false positive rate (Figure 5; median false positive rate of 0.22 [IQR: 0.17, 0.34]). As described above, the Inner Tukey observed a lower median false positive rate of 0.09 (IQR: 0.06, 0.18). In summary, both of our leave-one-out methods (Inner Tukey and MAD) produced a high true positive rate for rare variants with MAF > 0.001 (comparable to ADA) while maintaining low false positive rates across the entire spectrum of MAF.

**Figure 4.**
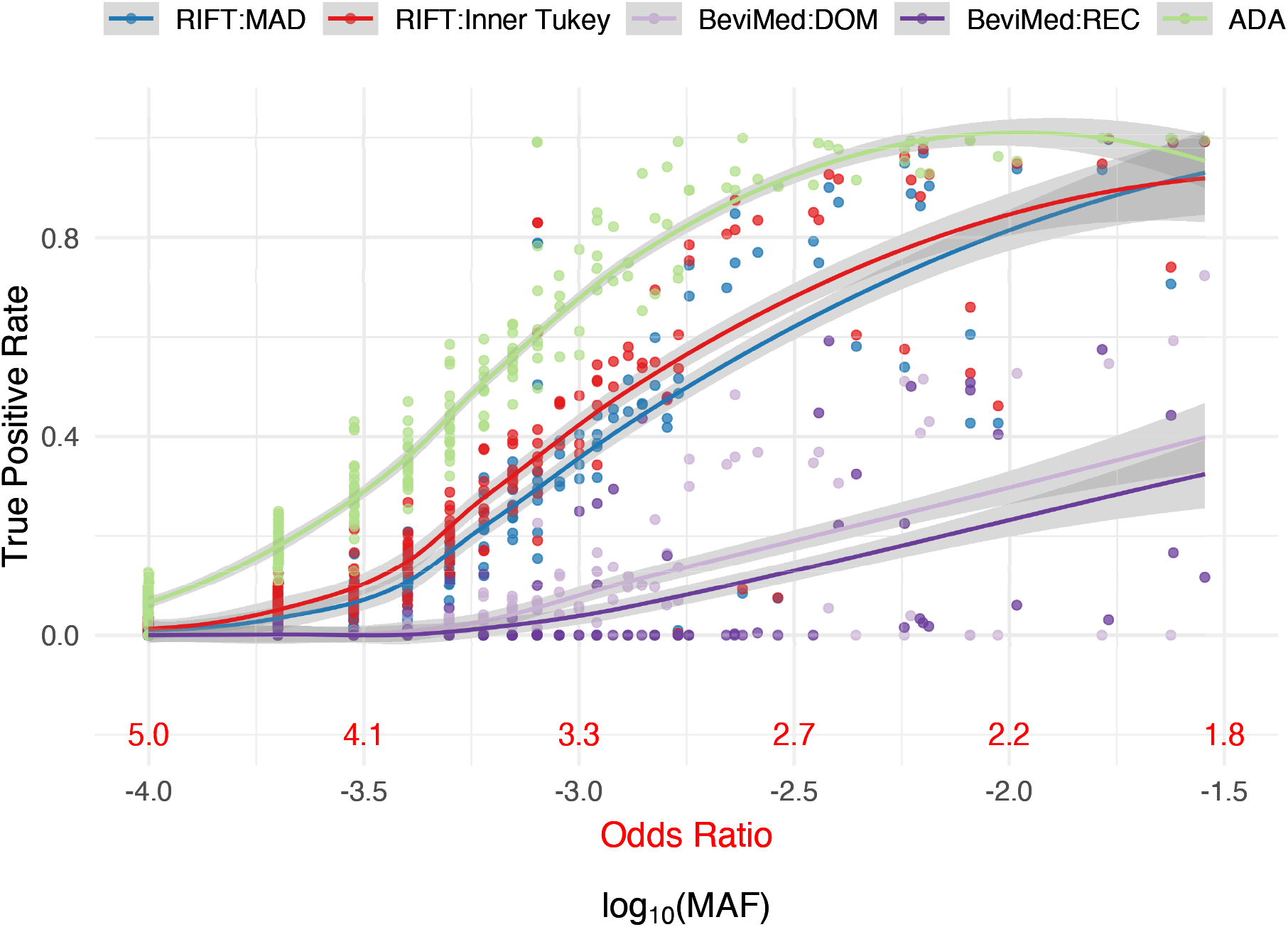
True positive rate (proportion correctly labeled IV) for variants under the alternative for each localization method as a function of MAF. Corresponding odds ratio is also provided for reference. Smoothed line and confidence band provided by the loess method. Data were simulated to have 10% variants under the alternative with effect size parameter *c* = 0.4 (see Equation 5).

**Figure 5.**
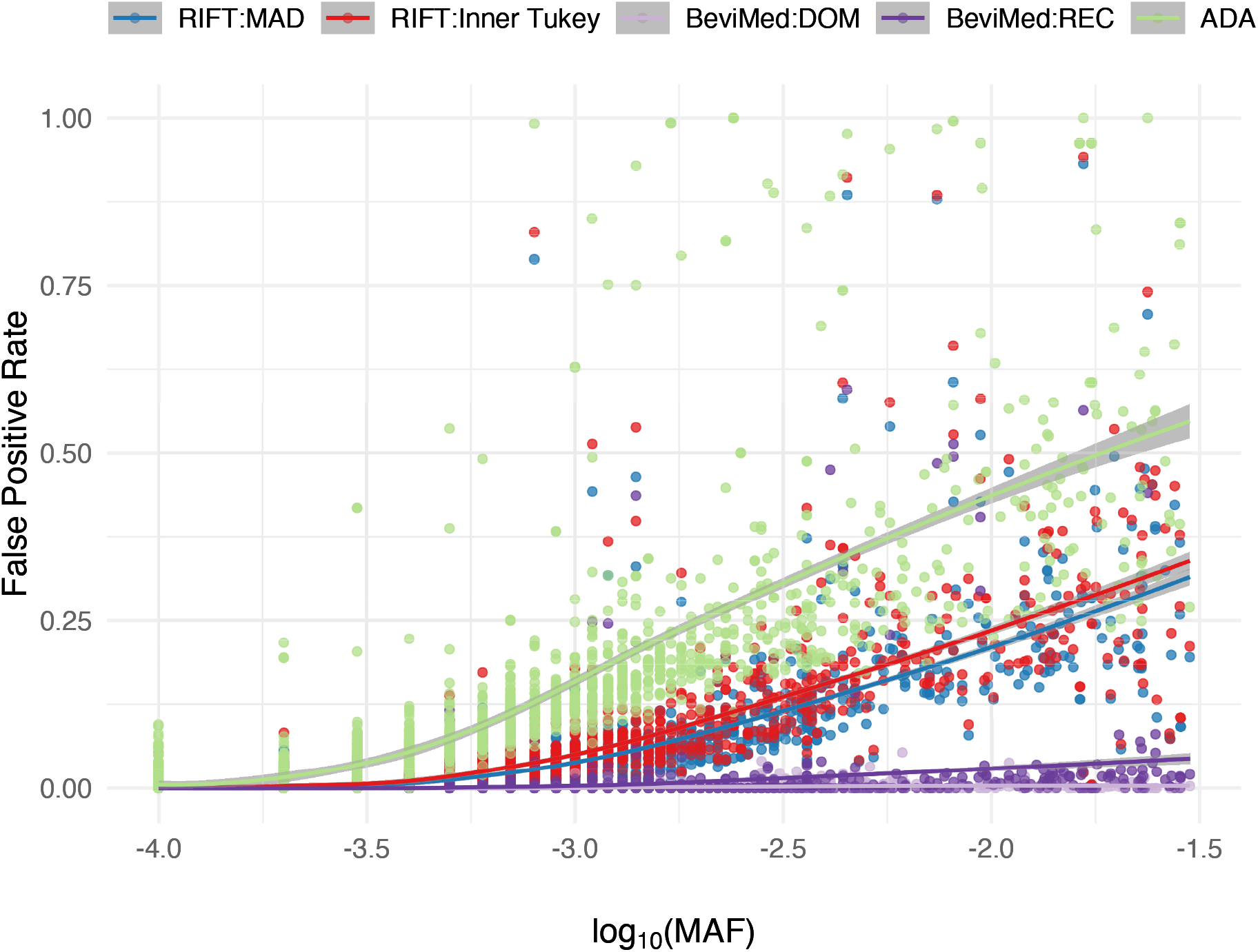
False positive rate (proportion incorrectly labeled IV) for variants under the null for each localization method as a function of MAF. Smoothed line and confidence band provided by the loess method. Data were simulated to have 10% variants under the alternative with effect size parameter *c* = 0.4 (see Equation 5).

### Influential Variants in IPF Associated Loci

In a rare variant analysis of targeted sequencing in 3,017 idiopathic pulmonary fibrosis (IPF) cases and 4,093 controls, we found several sets of rare variants associated with IPF. Rare variants were grouped into gene-sets or sliding windows and tested for association using SKAT-O. We applied RIFT to two significant rare variant windows, each of which contained 50 rare variants. The most significant window is located on chromosome 5 and spans the 5’ UTR, exon 1 and intronic regions of the *TERT* gene. This window had a Bonferroni-adjusted SKAT-O p-value of 9.21x 10^−16^ after adjusting for sex and the most strongly association common variant in the region, rs4449583. After applying RIFT, there were three variants called outliers by both the Inner Tukey fence and MAD cutoffs (Figure 6). Annotation with SNPDOC found the variant with the largest delta chi-square (−28.7) to have unknown function and the second ranked IV (delta chi-square = -17.5) to be in a non-coding RNA transcript in the 5’ untranslated region of *TERT* (Table 2). We additionally applied the localization method to a window with a less significant association (SKAT-O Bonferroni-adjusted p-value = 0.0215; adjusted for sex only as there is not a top common variant in the region) as an example of a region with a more moderate aggregate rare variant association signal. This window is located in the *RTEL* gene on chromosome 20 spanning both exons and introns. Our localization method identified 3 IVs by both the Inner Tukey fence and MAD cutoffs with the most outlying variant having a negative delta chi-square score = -6.77 (Figure 7; Table 2). This variant was annotated to be located in an intron of the *RTEL* gene and based on annotation with HaploReg, has enhancer histone marks identified in 8 tissues including lung and a lung carcinoma cell line. The other two IVs were each annotated to be in the coding region of *RTEL* and are nonsense mutations.

**Figure 6.**
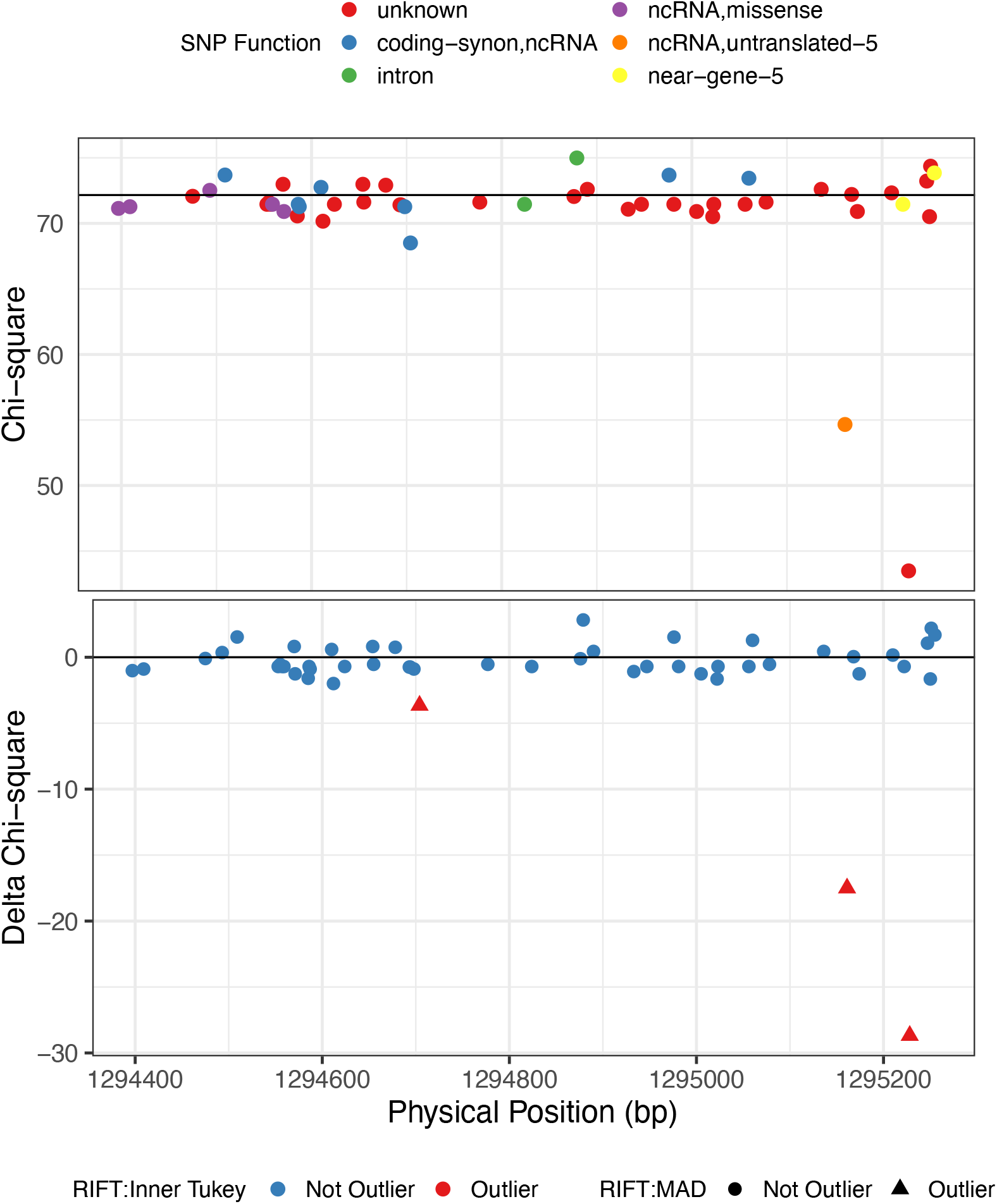
Chi-square (top and delta chi-square scores (bottom) by genomic position for the IPF-associated rare variant loci on chr5 (bp: 1294397-1295255). Color for the top plot corresponds to SNP-DOC functional annotation and for the bottom plot, color corresponds to outlier by the Inner Tukey method and shape corresponds to outlier by the MAD method.

**Figure 7.**
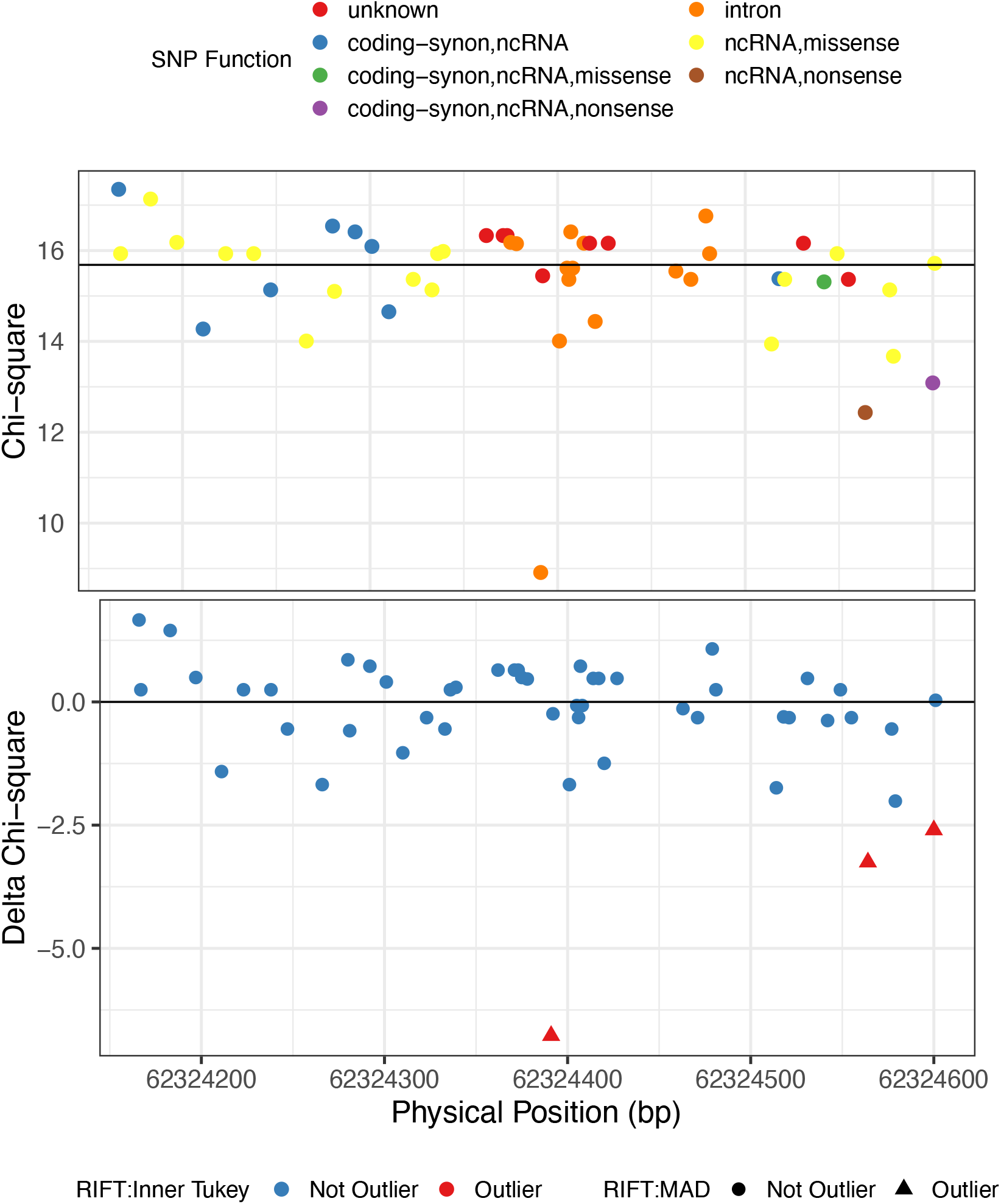
Chi-square (top and delta chi-square scores (bottom) by genomic position for the IPF-associated rare variant loci on chr20 (bp: 62324166-62324601). Color for the top plot corresponds to SNP-DOC functional annotation and for the bottom plot, color corresponds to outlier by the Inner Tukey method and shape corresponds to outlier by the MAD method.

## Discussion

The delta chi-square score we outline here provides an estimate of the contribution of a given variant to the aggregate test statistic for a set of variants. This measure can be used to rank variants in order to prioritize follow-up studies. We also compared several outlier detection methods to identify variants as having a disproportionate impact on the aggregate test of association, likely increasing the probability of capturing a causal variant. We found the inner Tukey fence to have the greatest sensitivity, and we recommend this method to obtain a set of variants most likely to be driving the aggregate signal. As expected, our method has higher sensitivity to detect uncommon variants (0.001 < MAF > 0.03) compared to extremely rare variants (MAF < 0.001). This underscores the difficulty in detecting extremely rare variants individually and thoughtful alternative weighting schemes might provide leverage to better capture very rare variants. The ADA method obtained the highest sensitivity for uncommon variants; however, this was at a cost of low specificity. We found BeviMed lacked sufficient sensitivity in the classification of IVs under our simulation framework. We recognize this may be due to using their recommended parameter of 0.90 posterior probability for classification of IVs. However, it is unclear how to determine an optimal value for the posterior probability, which likely depends on the features of each region (e.g., number of true causal variants, effect size of the variants, total number of variants tested). Without requiring user optimization or selection of parameters, RIFT with the Inner Tukey classification approach achieves higher sensitivity and specificity to ADA and BeviMed, respectively.

The rare variants identified by our method to have the greatest evidence for influencing the association with IPF have been previously found to be associated with other diseases (Table 2). Common, non-coding variants in the promoter region of the *TERT* gene have been found in several human cancers [29,30]. Specifically, the two IVs with the largest delta chi-square score (rs398123017 and rs373740199) have been identified in studies of familial and sporadic melanoma and thyroid cancer [31–34]. These mutations have been found to increase transcription of *TERT* and maintain telomerase activity resulting in long telomeres, thereby promoting tumorigenesis [35].

On chromosome 20, two of the three rare variants identified by our method are missense mutations in the *RTEL* gene and have been previously identified in dyskeratosis congenita and the related Hoyeraal-Hreidarsson syndrome, both of which are diseases that result due to failures in telomere maintenance [36–38]. Though extensive research was not performed for all variants within our significant sets (as we’ve previously noted is often prohibitive), identification of these variants in other diseases supports the ability of our approach to detect plausible impactful rare variants that drive our observed association with IPF.

Though we have only evaluated the delta chi-square statistic and corresponding outlier detection methods for data containing unrelated individuals, RIFT is not restricted to aggregate tests for this specific study design. Although we illustrated our approach using SKAT-O in a case-control framework, the methods are widely applicable to other aggregate tests and study designs. For example, the method could be applied using tests developed for samples of related individuals such as famSKAT [39].

Unlike many other localization methods, RIFT does not rely on outside functional information and is able to distinguish among variants with no known function that may be contributing to disease risk. We recognize that incorporating functional information can increase the power to identify causal rare variants. However, methods which rely on functional information are limited by the depth and accuracy of the annotation. Our method complements methods that utilize functional information to provide increased coverage in capturing plausible causal rare variation. Outlier detection methods to identify influential variants will eventually break down if a large proportion of the variants are under the alternative. It is plausible that in certain situations, the proportion of variants that are under the alternative is higher for a set of variants that all have putative function compared to a set that is selected agnostic of function. Since it is unlikely that the majority of putatively functional variants in a gene are associated with a given trait, it is also unlikely that filtering to functional variants prior to testing with an aggregate test would be problematic for the outlier detection methods implemented in RIFT. While RIFT maintains and perhaps even improves sensitivity for smaller numbers of variants included in the aggregate test (Supplemental Table 5), pre-filtering variants after the aggregate test is likely to be counter-productive. We recommend not filtering to variants with putative function either before or after aggregate testing, but instead applying RIFT agnostic of putative function and, if desired, further prioritizing those identified IVs based on putative function.

RIFT is available as an R package from https://github.com/rachelzoeb/RIFT. The time to complete an analysis with RIFT is dependent on the computation time of the aggregate test used. SKAT-O is fast for sets with a relatively small number of variants (e.g., < 100 rare variants) and requires parallelization for large significant sets. Future work will require optimization of RIFT for larger groups of variants such as sets based on gene boundaries.

Greater resolution of rare variant signals within significant sets of variants can provide valuable insight into the mechanism of disease and narrow the number of variants to prioritize for follow-up experimental studies. Often, aggregate tests contain such a large number of rare variants that follow up of each rare variant within the set would be prohibitive, even after filtering down to variants with functional evidence for causality. In addition, filtering to variants with known function precludes the ability to identify new variants with as-yet unknown functional roles. In addition to being agnostic to functional information, RIFT can be applied after application of any aggregate test, including those that include common variants. The ease and flexibility of this approach will aid investigators in post-aggregate association testing in a wide range of genetic studies.

## ACKNOWLEDGEMENTS

We acknowledge the Global IPF Collaborative Network (http://www.ucdenver.edu/academics/colleges/medicalschool/departments/medicine/GlobalIPF/Pages/GlobalIPF.aspx), the members of which contributed many of the DNA samples used to generate the resequencing data that are included in the example.

## STATEMENT OF ETHICS

For the IPF resequencing data, all of the subjects provided written informed consent as part of institutional review board (IRB)- approved protocols for recruitment at their respective institution, and the resequencing study was approved by the National Jewish Health IRB and the University of Colorado Combined IRB.

## AUTHOR’S CONTRIBUTIONS

CDL, TEF, and RZB designed the study and developed the conceptual approaches to data analysis; TEF and DAS provided resequencing data. RZB performed the simulations and data analysis; RZB wrote the manuscript; TEF, DAS, and CDL reviewed the manuscript.

