## Supplemental Material for "Identification of Influential Variants in Significant Aggregate Rare Variant Tests"

### SUPPLEMENTAL TABLES

**Supplemental Table 1. Summary of simulation parameters.** Values in bold correspond to the primary simulation conditions used to compare the various localization methods. Alternate values are explored and their impact on the resulting genotype data and performance of RIFT are described.

| Simulation Parameter | Value | Impact on simulations |
|---|---|---|
| # haplotypes simulated | 1,000 / **10,000** / 100,000 | - Decreased sampling of haplotypes for 3kb region results in smaller number of total variants and reduces the power of SKAT-O (ST3).<br>- Number of variants does not impact RIFT (ST5). |
| Region size (kb) | 0.75 / **3** | - Smaller region size results in smaller number of total variants and reduces the power of SKAT-O (data not shown).<br>- Number of variants does not impact RIFT (ST5) |
| # cases/controls | **1000** / 5000 | - Increasing the sample size results in increased power of SKAT-O (ST3) and larger negative delta chi-square scores for variants under the alternative (SF2) |
| Disease prevalence | **0.05** / 0.10 | - Results similar between 0.05 and 0.10 prevalence (data not shown) |
| Coefficient of disease association *c* | **0.4** / 0.8 | - Increasing the coefficient of disease association increased the power of SKAT-O (ST2) and improves ability of RIFT to detect influential variants (data not shown) |
| Proportion alternative variants | **0.10 /** 0.20 | - Increasing the proportion variants under the alternative increased the power of SKAT-O (ST2) and improves ability of RIFT to detect influential variants (data not shown) |

**Supplemental Table 2. Summary of simulations varying proportion variants under the alternative and effect size**. Each simulation consisted of 50 regions with 1000 samples per region. Results in manuscript reported for simulations with 10% proportion variants under the alternative and effect size parameter c = 0.4 (see Equation 5).

| % alternative simulated | *c* | Average # total variants per region Median (IQR) | Average % alternative per region Median (IQR) | # samples SKAT-O p < 0.05 (out of 1000) Median (IQR) |
|---|---|---|---|---|
| 0.10 | 0.40 | 39.56 (35.43, 43.14) | 0.12 (0.12, 0.13) | 220 (109, 520.25) |
| 0.10 | 0.80 | 40.35 (36.42, 43.70) | 0.14 (0.13, 0.15) | 990.50 (764.50, 1,000) |
| 0.20 | 0.40 | 38.37 (35.61, 41.94) | 0.24 (0.23, 0.26) | 727 (427.25, 947.50) |
| 0.20 | 0.80 | 39.26 (36.50, 43.04) | 0.27 (0.25, 0.28) | 1000 (1000, 1000) |

**Supplemental Table 3. Summary of simulations varying sample size and number of haplotypes simulated**. Each simulation consisted of 10 regions with 500 samples per region. Results in manuscript reported for simulations with 10% proportion variants under the alternative and effect size parameter c = 0.4 (see Equation 5).

| # of haplotypes simulated | # Case/ # Control | Average # total variants per region Median (IQR) | Average % alternative per region Median (IQR) | # SKAT-O samples < 0.05 (out of 500) Median (IQR) |
|---|---|---|---|---|
| **1k** | 1000/1000 | 20 (18.00, 21.00) | 2 (2.00, 2.00) | 117.0 (64.5, 318.3) |
| **10k** | 1000/1000 | 60 (54.00, 64.00) | 6 (5.00, 6.00) | 222.0 (106.5, 413.0) |
| **10k** | 5000/5000 | 60.00 (56.00, 67.75) | 6 (6.00, 7.00) | 457.5 (373.8, 495.5) |
| **100k** | 1000/1000 | 484 (475.00, 512.00) | 48 (48.00, 51.00) | 272.0 (191.8, 361.8) |

**Supplemental Table 4.** Summary of true positive rate (TPR) and false positive rate (FPR) of the four outlier detection methods separated by MAF < 0.001 and MAF ≥ 0.001.

| Method | True Positive Rate - Median (IQR) | | False Positive Rate – Median (IQR) | |
|---|---|---|---|---|
| | MAF < 0.001 | MAF ≥ 0.001 | MAF < 0.001 | MAF ≥ 0.001 |
| **RIFT:MAD** | 0.03 (0.01, 0.11) | 0.54 (0.43, 0.85) | 0.00 (0.00, 0.01) | 0.08 (0.04, 0.15) |
| **RIFT:SD** | 0.00 (0.00, 0.00) | 0.30 (0.10, 0.59) | 0.00 (0.00, 0.00) | 0.00 (0.00, 0.04) |
| **RIFT:Inner Tukey** | 0.04 (0.01, 0.15) | 0.60 (0.51, 0.87) | 0.00 (0.00, 0.01) | 0.09 (0.06, 0.18) |
| **RIFT:Outer Tukey** | 0.01 (0.00, 0.05) | 0.40 (0.30, 0.81) | 0.00 (0.00, 0.00) | 0.05 (0.02, 0.11) |

**Supplemental Table 5.** Summary of true positive rate (TPR) and false positive rate (FPR) of the four outlier detection methods separated by MAF < 0.001 and MAF ≥ 0.001 for simulations of smaller regions (0.75kb).

| Method | True Positive Rate - Median (IQR) | | False Positive Rate – Median (IQR) | |
|---|---|---|---|---|
| | MAF < 0.001 | MAF ≥ 0.001 | MAF < 0.001 | MAF ≥ 0.001 |
| **RIFT:MAD** | 0.07 (0.00, 0.21) | 0.79 (0.58, 0.89) | 0.00 (0.00, 0.01) | 0.13 (0.06, 0.27) |
| **RIFT:SD** | 0.00 (0.00, 0.03) | 0.04 (0.02, 0.18) | 0.00 (0.00, 0.00) | 0.00 (0.00, 0.06) |
| **RIFT:Inner Tukey** | 0.05 (0.00, 0.19) | 0.79 (0.64, 0.85) | 0.00 (0.00, 0.00) | 0.13 (0.06, 0.29) |
| **RIFT:Outer Tukey** | 0.00 (0.00, 0.09) | 0.64 (0.42, 0.73) | 0.00 (0.00, 0.00) | 0.09 (0.03, 0.21) |

To show that RIFT remains valid for a smaller number of rare variants included in the aggregate test of association (and therefore included in the RIFT procedure), we simulated genetic regions of a smaller size to achieve an average of 10 variants per region. Using the same approach we used with 3kb regions, we simulated 50 750bp regions with 10% of variants under the alternative. After applying RIFT to only samples which met SKAT-O significance, the resulting samples observed an average of 9.8 variants with 1.3 variants under the alternative. The median TPR of the Inner Tukey outlier detection method among uncommon variants (MAF ≥ 0.001) was 0.79 (IQR: 0.64, 0.85), higher for the smaller regions compared to what was observed for the 3kb regions (Supplemental Table 5). The median TPR of 0.05 (IQR: 0.00, 0.19) among very rare variants (MAF < 0.001) of the 0.75kb regions was comparable to the 3kb regions. Thusly, RIFT does not require a minimum number of variants to prioritize influential variants efficiently.

### SUPPLEMENTAL FIGURES

**Supplemental Figure 1. Proportion called outliers for variants under the null by outlier detection methods**. Overall, proportion of null variants labeled IVs is low across all methods and is highest for variants having larger MAF. The parametric approach of 3 standard deviations performs best by having the lowest proportion of variants inaccurately identified as IVs.

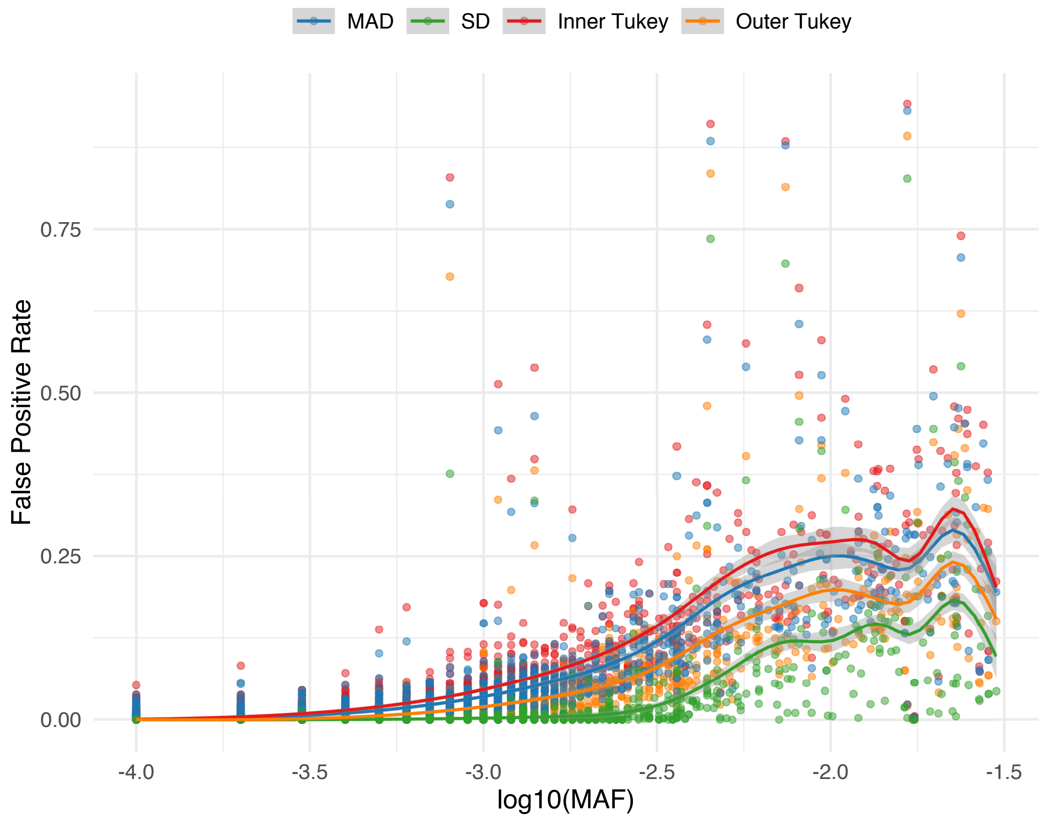

**Supplemental Figure 2. Relationship between delta chi-square score and MAF stronger for larger sample size and consistent with varying number of simulated haplotypes.** Mean delta chi-square score plotted by MAF for variants simulated under the alternative across 10 regions. Data were simulated to have 10% variants under the alternative with effect size parameter *c* = 0.4 (see Equation 5).

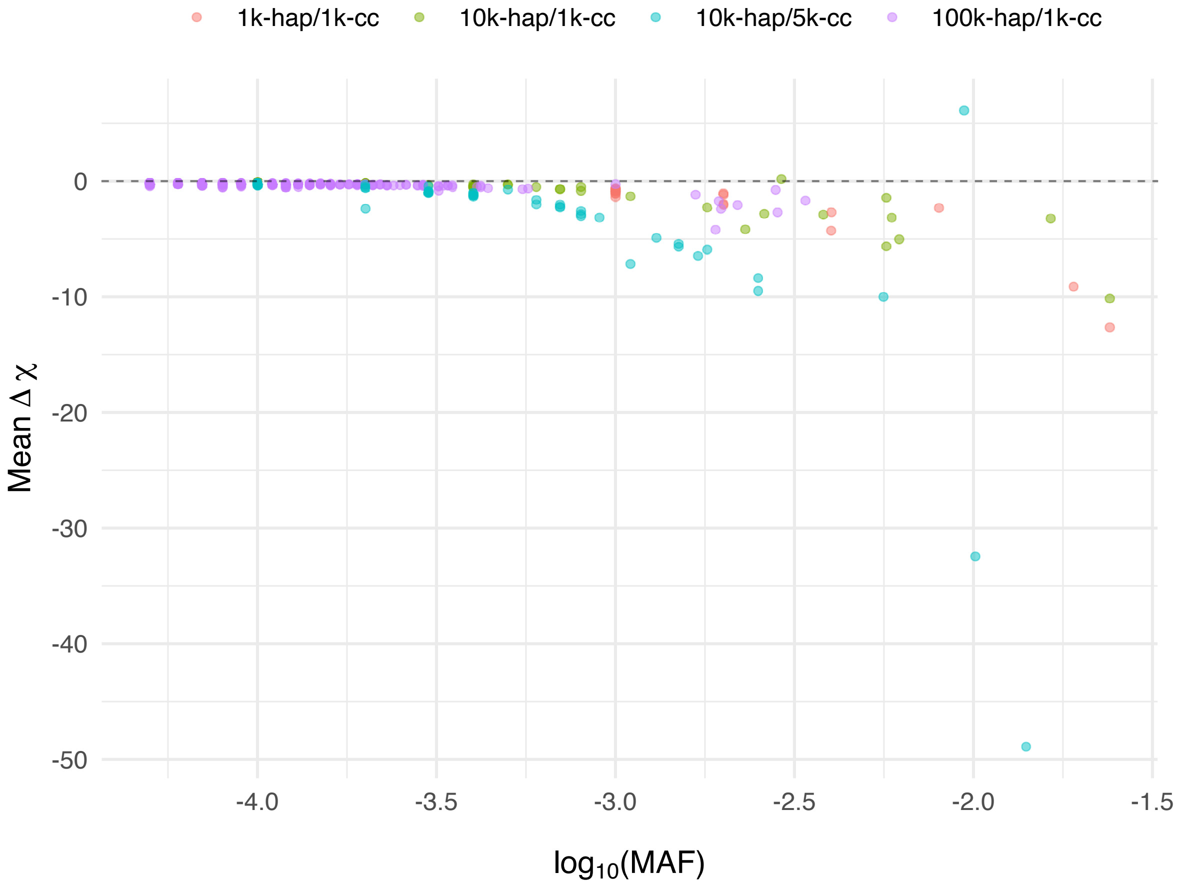
